# A framework for spatial normalization and voxelwise analysis of diffusion propagators in multiple MAP-MRI data sets

**DOI:** 10.1101/697284

**Authors:** Alexandru V. Avram, Adam S. Bernstein, M. Okan Irfanoglu, Craig C. Weinkauf, Martin Cota, Neville Gai, Amber Simmons, Anita Moses, L. Christine Turtzo, Neekita Jikaria, Lawrence Latour, Dzung L. Pham, John A. Butman, Peter J. Basser

**Affiliations:** National Institute of Biomedical Imaging and Bioengineering, National Institutes of Health, Bethesda, MD, 20892, USA; Eunice Kennedy Shriver National Institute of Child Health and Human Development, National Institutes of Health, Bethesda, MD, 20892, USA; University of Arizona, Department of Biomedical Engineering, Tucson, AZ, 85724, USA; University of Arizona, Department of Surgery, Tucson, AZ, 85724, USA; Center for Neuroscience and Regenerative Medicine, Henry Jackson Foundation, Bethesda, MD, 20892, USA; Clinical Center, National Institutes of Health, Bethesda, MD, 20892, USA; National Institute of Neurological Disorders and Stroke, National Institutes of Health, Bethesda, MD, 20892, USA

**Keywords:** MAP-MRI template, propagator template, diffeomorphic registration, tensor-based image registration, tissue microstructure, diffusion tensor imaging (DTI)

## Abstract

We describe a pipeline for constructing a study-specific template of diffusion propagators measured with mean apparent propagator (MAP) MRI that supports direct voxelwise analysis of differences between propagators across multiple data sets. The pipeline leverages the fact that MAP-MRI is a generalization of diffusion tensor imaging (DTI) and combines simple and robust processing steps from existing tensor-based image registration methods. First, we compute a DTI study template which provides the reference frame and scaling parameters needed to construct a standardized set of MAP-MRI basis functions at each voxel in template space. Next, we transform each subjects diffusion data, including diffusion weighted images (DWIs) and gradient directions, from native to template space using the corresponding tensor-based deformation fields. Finally, we fit MAP coefficients in template space to the transformed DWIs of each subject using the standardized template of MAP basis functions. The consistency of MAP basis functions across all data sets in template space allows us to: 1. compute a template of propagators by directly averaging MAP coefficients and 2. quantify voxelwise differences between co-registered propagators using the angular dissimilarity, or a probability distance metric, such as the Jensen-Shannon Divergence. We illustrate the application of this method by generating a template of MAP propagators for a cohort of healthy volunteers and show a proof-of-principle example of how this pipeline may be used to detect subtle differences between propagators in a single-subject longitudinal clinical data set. The ability to standardize and analyze multiple clinical MAP-MRI data sets could improve assessments in cross-sectional and single-subject longitudinal clinical studies seeking to detect subtle microstructural changes, such as those occurring in mild traumatic brain injury (mTBI), or during the early stages of neurodegenerative diseases, or cancer.

## 1. Introduction

Diffusion MRI (dMRI) is a clinical tool that probes non-invasively the motion of water molecules in tissues and can thereby be very sensitive to alterations in tissue microstructure and cytoarchitectural organization during disease. In general, dMRI signals in both isotropic and anisotropic tissues can be accurately described with a diffusion tensor model (Basser et al., 1994). Even though diffusion tensor imaging (DTI) has been shown to be sensitive to brain tissue changes in stroke, cancer and other diseases, there is a need to improve the sensitivity of dMRI analysis for detection of subtle tissue changes, such as those occurring during mild traumatic brain injury (mTBI) or during the early stages of neurodegenerative diseases. Two avenues to improve the sensitivity of dMRI are by: 1. analyzing diffusion weighted images (DWIs) acquired with high diffusion sensitization (b-value) and 2. analyzing multiple standardized data sets in cross-sectional or longitudinal clinical studies.

While the DTI model can explain most of the brain dMRI signal variations, it doesn’t accurately capture the signal modulations occurring in tissue regions with crossing white matter (WM) fiber pathways at high b-values, and requires refinement using higher-order terms. A comprehensive dMRI assessment of tissue microstructure can be obtained by explicitly measuring the 3D probability distributions of microscopic net displacements of water molecules in tissues, i.e., the diffusion propagators, from DWIs acquired with a wide range of b-values and many gradient orientations. Mean apparent propagator (MAP) MRI (Özarslan et al., 2013) provides an efficient, elegant, and clinically feasible (Avram et al., 2016) solution that subsumes and analytically extends the DTI model. In every voxel, the dMRI signal is approximated using MAP-MRI basis functions specifically defined in the reference frame determined by the principal axes of the diffusion tensor, using the principal diffusivities as scaling parameters (Özarslan et al., 2013). This formulation provides a compact and efficient analytical approximation of the propagators from only the first few terms (i.e., coefficients) in the MAP-MRI series expansion and allows direct comparison of new clinical findings with results from the existing DTI literature (Avram et al., 2016).

The ability to conduct voxel-wise analysis of dMRI signals across multiple co-registered MAP data sets could further strengthen the sensitivity of detecting and characterizing subtle pathology-related tissue changes in cross-sectional studies using test and control groups, or in longitudinal studies comparing tissue properties at different time-points within the same subject. Nevertheless, both cross-sectional and single-subject longitudinal studies require adequate tools for accurate spatial normalization and analysis of multiple MAP-MRI data sets. Such tools would also enable the creation of MAP-MRI templates that may serve as normative databases (Zhang and Arfanakis, 2018) for pothole analysis (Watts et al., 2014; White et al., 2009) in studying pathologies with subtle and diffuse presentations such as mTBI or the early stages of neurodegenerative diseases. They could also improve the identification, delineation, labelling and characterization of isotropic and anisotropic anatomical structures in brain atlases (Mori et al., 2009) based on differences in their tissue microstructural and cytoarchitectural properties.

Many scalar and tensor-based algorithms for diffeomorphic registration of multiple DTI data sets (Alexander et al., 2001; Jones et al., 2002; Gee and Alexander, 2006; Zhang et al., 2006; Muoz-Moreno et al., 2009) and are used in a growing number of developmental and clinical studies (Sadeghi et al., 2015; Maier-Hein et al., 2015; Poudel et al., 2015; Mahoney et al., 2015; Garaci et al., 2015; White et al., 2009; Wang et al., 2011). More recently, a number of algorithms have been proposed to register dMRI data sets acquired with high b-values using: higher-order signal representations (Barmpoutis et al., 2007), orientation distribution functions (ODFs) (Geng et al., 2009), fiber ODFs (FODs) (Raffelt et al., 2011), propagators (Ginsburger et al., 2018) or by directly aligning DWIs (Chiang et al., 2008; Zhang et al., 2014). Many of these methods employ complicated new processing steps and multiparametric optimizations, and focus primarily on the orientational alignment in regions of crossing WM fibers to improve mapping of neural pathways and brain connectivity. On the other hand, tensor-based method, such as DRTA-MAS (Irfanoglu et al., 2016), provide a simple and robust solution for dMRI data standardization for analyzing a wide range of microstructural properties. When applied to DWIs, tensor-based transformation fields produce repeatable results and improve spatial alignment in all white matter, including regions of crossing fibers (Irfanoglu et al., 2016). For MAP-MRI analysis tensor-based diffeomorphic registration provides additional benefits. Since it works in Cartesian coordinates, it is directly compatible with DTI and MAP-MRI and does not require new complicated analytical operations and processing steps, such as change of basis transformations between signal representations, which can lead to truncation errors. More importantly, tensor-based diffeomorphic registration inherently provides the DTI study template necessary for voxelwise analysis of co-registered propagators using a common template of MAP basis functions.

In this study, we describe an efficient and robust pipeline for spatial normalization of multiple MAP-MRI data sets and voxelwise analysis across co-registered propagators. The proposed method can provide maps of angular dissimilarity between the propagators of a single subject and the template propagators. The ability to quantify voxelwise differences between propagators may improve the assessment of subtle microstructural features in cross-sectional group and single-subject longitudinal clinical studies.

## 2. Methods

### 2.1. MAP-MRI data acquisition from multiple subjects

We scanned twelve healthy volunteers with no documented history of traumatic brain injury (TBI) and no radiological findings on a clinical 3T GE MR750 MRI scanner. All subjects participating in this study provided written and informed consent in accordance with a clinical protocol approved by the institutional review board (IRB) of the Intramural Research Program (IRP) at the NIH Clinical Center and the Center of Neuroscience and Regenerative Medicine (CNRM). We acquired MAP DWIs using a spin-echo diffusion echo-planar imaging (EPI) pulse sequence with the following imaging parameters: TE/TR=94/6000ms, 42 slices with 3mm slice thickness on a 210 × 210 *mm*^2^ field-of-view (FOV) with a 70 × 70 imaging matrix, SENSE factor of 2, and no partial Fourier sampling, resulting in a 3mm isotropic voxel resolution. A total of 498 DWIs were acquired with multiple b-values: 0, 1000, 2000, 3000, 4000, 5000, and 6000 *s/mm*^2^, with increasing number of diffusion encoding gradient directions at larger b-values. The gradient orientations were chosen to uniformly sample the unit sphere at each b-value and across b-shells (Cheng et al., 2014). The diffusion gradient pulse width and separation were *δ* = 34.2*ms* and ∆ = 40.2*ms*, respectively. To quantify the signal-to-noise ratio (SNR) and estimate a field map for subsequent correction for EPI distortions due to *B*_0_ magnetic field inhomogeneities, we obtained an additional series of 7 consecutive images with reversed EPI phase encoding and identical imaging parameters, but no diffusion sensitization. Additionally, we acquired a high resolution fast spin echo (FSE) T2-weighted image to serve as an anatomical template for image registration and motion correction of DWIs, using a 1mm spatial resolution, an FOV of 240mm, TE/TR=245/2000ms, and 240 echoes; and a T1-weighted MP-RAGE scan for automatic segmentation and parcellation in the ROI analysis with 1mm spatial resolution, an FOV of 240mm, TE/TI/TR=92.3/1703/5000ms.

We processed all MAP DWIs using the TORTOISE software package (Pierpaoli et al., 2010) to correct for Gibbs ringing correction (Kellner et al., 2016), subject motion correction and eddy current induced EPI distortions. DWI denoising using principal component analysis as described in (Veraart et al., 2015) was performed as an intermediate step to improve the image registration steps, without propagation to the final preprocessed image. Eddy current distortions were corrected by registering each DWI to the anatomical T2-weighted reference image using quadratic deformations. The phase-reversed scan was processed similarly with TORTOISE to correct for subject motion. Subsequently, we combined the two motion and eddy current corrected data sets (blip-up and blip-down) with DRBUDDI (Irfanoglu et al., 2015) to correct for additional EPI distortions due to static magnetic field inhomogeneities. Finally, the DWIs were interpolated from 3mm to 2mm isotropic resolution.

### 2.2. MAP-MRI analysis in subject space

We analyzed the diffusion data sets in native (subject) space by fitting MAP-MRI coefficients (Özarslan et al., 2013) on a voxel-by-voxel basis as described in (Avram et al., 2016). First, we estimated a diffusion tensor using weighted linear least squares from all DWIs with b-values ≤ 2000*s/mm*^2^. Then, we used the orientation and magnitudes of the principal diffusivities of the diffusion tensors to construct the set of MAP-MRI basis functions in each voxel (Appendix A). We computed the MAP-MRI coefficients up to order 6 using quadratic programming with constraints to ensure the propagator was normalized and positive for all displacements. From the MAP coefficients we derived microstructural MAP parameter maps of return-to-origin probability (*RTOP*), return-to-axis probability (*RTAP*), return-to-plane probability (*RTPP*), propagator anisotropy (*PA*), gaussian propagator anisotropy (*PA*_*DTI*_), non-gaussian diffusion anisotropy (∆Θ_*PO*_), non-gaussianity (*NG*), axial non-gaussianity (*NG*_||_), and radial non-gaussianity (*NG*_⊥_) (Avram et al., 2016).

### 2.3. Constructing the MAP study template

The motion and distortion corrected DWIs from all subjects were processed to construct a study-specific MAP template using simple processing steps as shown in Fig. 1:

**Figure 1:**
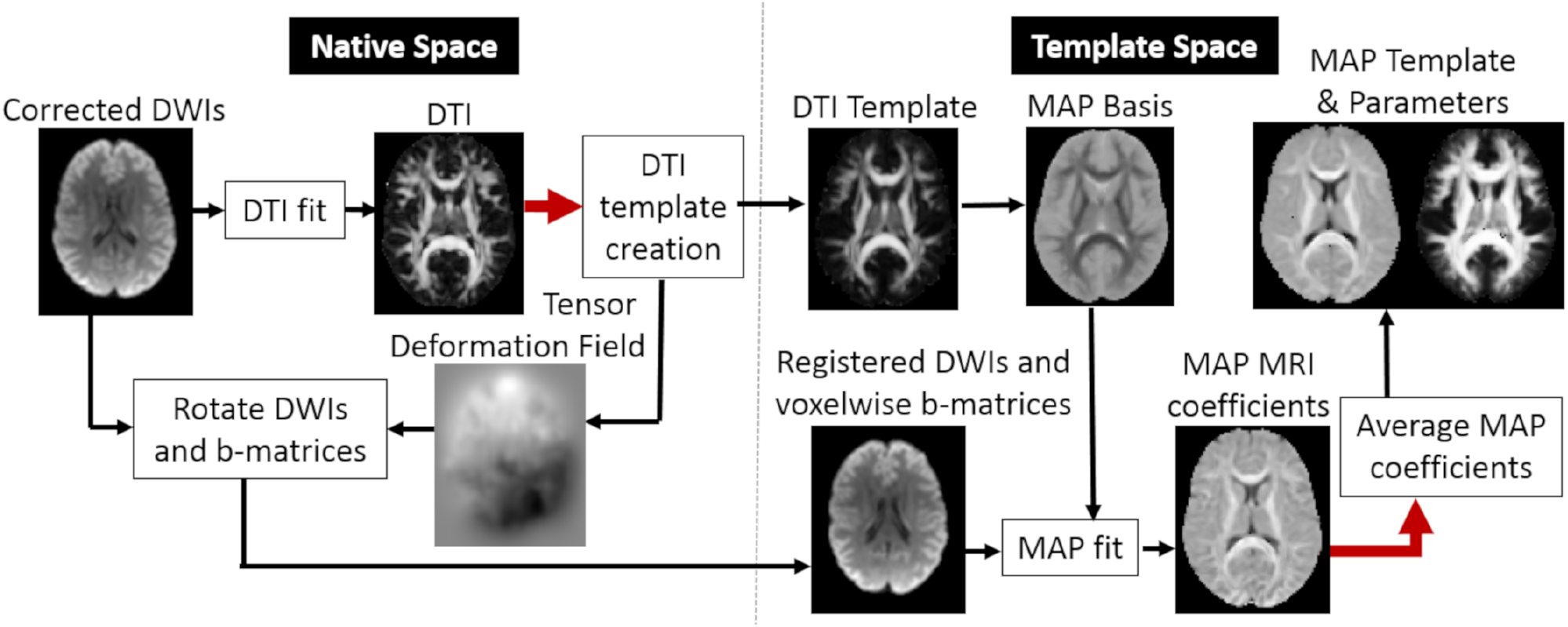
A flowchart for spatial standardization of MAP-MRI data and for constructing a population template of MAP-MRI diffusion propagators.

- To start with, we generated a DTI study template by registering the subset of DWIs acquired with low b-values, ≤ 2000*s/mm*^2^, from each MAP data set using the tensor-based diffeomorphic registration framework of DRTA-MAS (Irfanoglu et al., 2016).
- Then, we applied the tensor-based deformation fields produced in the previous step to transform the DWIs and corresponding diffusion encoding b-matrices in each data set from native to template space, as described in (Irfanoglu et al., 2016).
- Next, we computed the eigenvectors and eigenvalues (principal diffusivities) from the DTI study template and used them as orientation and scaling parameters necessary to construct the MAP basis functions standardized for each voxel location in the template.
- In the end, we estimated the MAP series coefficients up to the sixth order in template space by fitting the transformed DWIs and corresponding voxel-wise b-matrices of each co-registered data set to the same standardized template of MAP basis functions derived from the DTI study template.

The last key step of the pipeline ensures that in each voxel we quantify the propagators in template space using a consistent set of MAP basis functions for all co-registered data sets. This allows us to voxelwise average and compare MAP coefficients, microstructural parameters, and even entire propagators.

For example, we computed a population template of MAP-MRI propagators by voxelwise averaging of MAP coefficients across subjects. Since the coefficients in each MAP data set represent positive and normalized propagators (i.e., proper probability density functions), averaging across data sets preserves these properties for the template propagators. From the template of 3D diffusion propagators we derived MAP microstructural parameters to compare with similar metrics quantified in individual subject space. We also computed orientation profiles of the template propagators, i.e., orientation distribution functions (ODFs) (Özarslan et al., 2013). We derived fiber ODFs (fODFs) using spherical deconvolution (Tournier et al., 2004, 2007) and visualized them using the MRTrix3 software package (Tournier et al., 2012). Finally, we performed whole-brain streamline fiber tractography using the template fODFs.

### 2.4. Voxel-wise comparison of propagators in co-registered MAP datasets

Using a voxel-wise standardized MAP functional basis to represent and analyze spatially normalized 3D diffusion propagators allows direct assessment of subtle microstructural differences across multiple subjects. For example, the angular dissimilarity proposed by (Özarslan et al., 2013) is a convenient way to quantify differences between two propagators expressed in the same MAP functional basis. The angular dissimilarity, sin *θ*, between two 3D MAP propagators expressed in the same orthogonal MAP functional basis using MAP coefficient column vectors **a** and **b**, respectively, can be computed as:

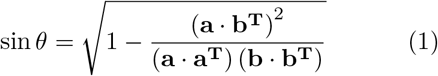

The angular dissimilarity is analogous to computing the angle between two vectors in Cartesian space using the dot product and underlies the computation of the PA and NG; the PA quantifies the angular dissimilarity between the MAP propagator and its isotropic approximation, while the NG quantifies the angular dissimilarity between the full propagator and its Gaussian (diffusion tensor) approximation. The proposed framework can provide maps of angular dissimilarity values between any two co-registered propagators in template space (for example single subject vs. population average). Such maps could be exquisitely sensitive to intersubject microstructural differences reflected in relevant propagator features that are not fully captured by anisotropy, diffusivity, or non-gaussianity measures independently.

### 2.5. Region-of-interest based analysis

To validate the performance of our pipeline, we conducted region-of-interest (ROI) based analysis of MAP-MRI parameters measured in subject space and template space, and compared the results. We identified white matter structures based on the International Consortium of Brain Mapping (ICBM)-DTI-81 atlas. In subject space, the ICBM-DTI-81 FA map was registered to each subject’s FA map, and the transformation was then applied to the parcellation map (atlas labels) using nearest-neighbor interpolation. The means and standard deviations of MAP microstructural parameters PA, NG, and RTOP were quantified in each registered ROI. In template space, the same procedure was followed using an FA map derived from the DTI population template to register the ROIs. We computed means and standard deviations for PA, NG, and RTOP maps of each subject in template space and compared them to the ROI statistics of the same parameters measured in subject space.

### 2.6. Longitudinal MAP-MRI data acquisition and analysis

To illustrate the potential clinical application of spatial standardization and analysis of MAP-MRI measurements we analyzed preliminary data from a single-subject longitudinal study. Specifically, we collected data from one subject with a history of severe unilateral asymptomatic carotid atherosclerosis (*>*70% occlusion) undergoing surgical treatment who provided written and informed consent in accordance with a clinical protocol approved by the IRB of the University of Arizona. MAP-MRI data was acquired at three timepoints using a 3T MRI scanner. The first baseline scan was performed one week prior to carotid endarterectomy. The second scan was performed one month following the procedure, and the third and final scan was performed six months following the procedure. At each timepoint, the MAP-MRI experiment used a spinecho EPI sequence with TR/TE = 10700/115 ms, a matrix size of 128 × 128 and 69 slices, an FOV (256 × 256 mm) with 2 mm isotropic resolution, an in-plane GRAPPA factor of 2, and 14 b = 0 *s/mm*^2^, 20 b = 1000 *s/mm*^2^, 32 b = 2000 *s/mm*^2^, and 64 b = 3000 *s/mm*^2^, with diffusion gradient orientations determined by a multishell electrostatic repulsion scheme similar to that described by (Caruyer et al., 2013), producing a direction set that is both evenly distributed on each shell and non-collinear across shells. In addition, a single b = 0 *s/mm*^2^ was collected with the phase encoding direction reversed for use in EPI distortion correction.

The MAP-MRI data sets acquired at the three different timepoints were processed using the spatial standardization pipeline described in Fig. 1. In short, we first generated a DTI template from the low b-value DWIs (b ≤ 2000*s/mm*^2^) at the three timepoints. Using the tensor-based deformation fields, the MAP DWIs and corresponding gradient orientations at each timepoint were transformed to template space, where they were analyzed using a MAP-MRI fit up to order 4 with basis functions determined by the DTI template. Microstructural changes were quantified in template space on a voxel-by-voxel basis using the angular dissimilarity measure (Eq. 1) between timepoints.

To quantify subtle differences between MAP propagators longitudinally we propose a new approach using the Jensen-Shannon Divergence (JSD). As shown in Appendix B, the JensenShannon divergence between two MAP propagators (i.e., 3D probability density functions of net displacements of diffusing water molecules), *P* (**r**) and *Q*(**r**), defined in the same functional basis by the column vectors of MAP coefficients **a** and **b** respectively, can be analytically approximated as:

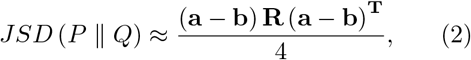

where **R** is a constant scalar matrix defined in Appendix B. We compute the voxel-wise JSD between the three timepoints in the longitudinal data set and compare it to the angular dissimilarity metric.

## 3. Results

The average brain tissue signal-to-noise ratio (SNR) in the non-diffusion weighted images in the MAP data sets was approximately 80, which is sufficient for computation of unbiased values of MAP parameters (Avram et al., 2016). After postprocessing using TORTOISE, all MAP DWIs, including those acquired with high b-values, showed good spatial agreement with the structural T2weighted scans, and little distortions due to eddy currents, *B*_0_ imhomogeneities, and/or subject motion. For all subjects, microstructural DTI and MAP parameters computed in the subject space were within the expected value ranges and showed the expected contrasts between brain tissues.

Microstructural MAP-MRI parameters derived from the template 3D diffusion propagators were consistent with previous clinical measurements in healthy brain tissue (Avram et al., 2014, 2016) (Fig. 2). Despite the 3mm acquisition resolution, and the anatomical variations across subjects, the RTOP, PA, and NG maps showed good delineation of white matter (WM) and gray matter (GM) tissues. Fiber ODFs derived from population-averaged template 3D propagators showed multiple peaks in regions of crossing white matter fiber bundles. We can resolve crossing WM pathways using whole-brain streamline fiber tractography through the template fODF field (Fig. 3B). It is also interesting to note that, to some extent, the orientational profiles show differences across GM regions (Fig. 3A).

**Figure 2:**
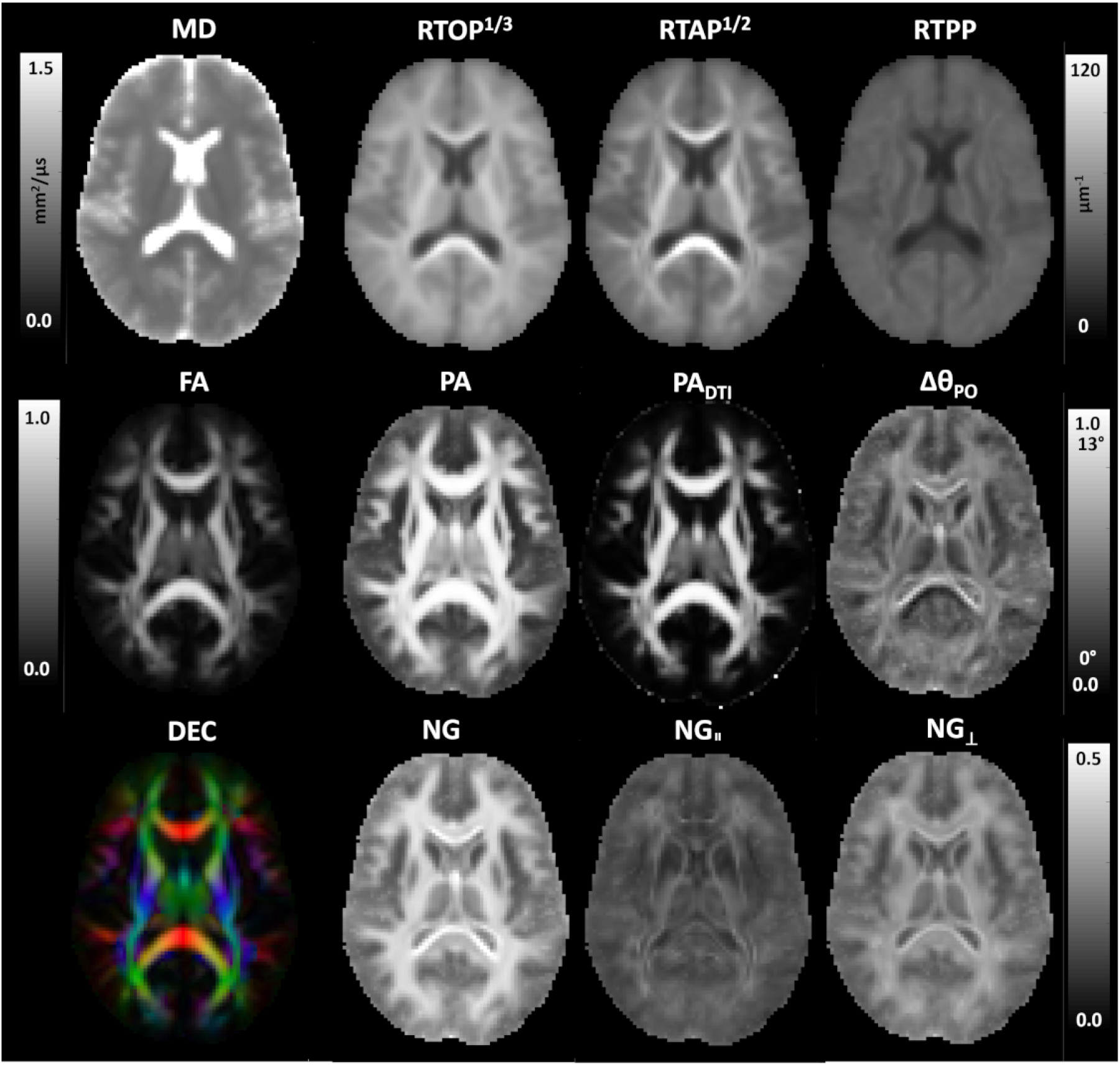
Microstructural MAP-MRI parameters computed from the template of diffusion propagators measured in the cohort of 12 healthy volunteers: DTI-parameters: mean diffusivity (MD), fractional anisotropy (FA), and direction encoded color (DEC) map; along with MAP microstructural parameters: return-to-origin probability (RTOP), return-to-axis probability (RTAP), return-to-plane probability (RTPP), propagator anisotropy (PA), gaussian propagator anisotropy (*PA*_*DTI*_), nongaussian anisotropy (∆Θ_*PO*_), non-gaussianity (NG), axial non-gaussianity (*NG*_||_), and radial non-gaussianity (*NG*_⊥_).

**Figure 3:**
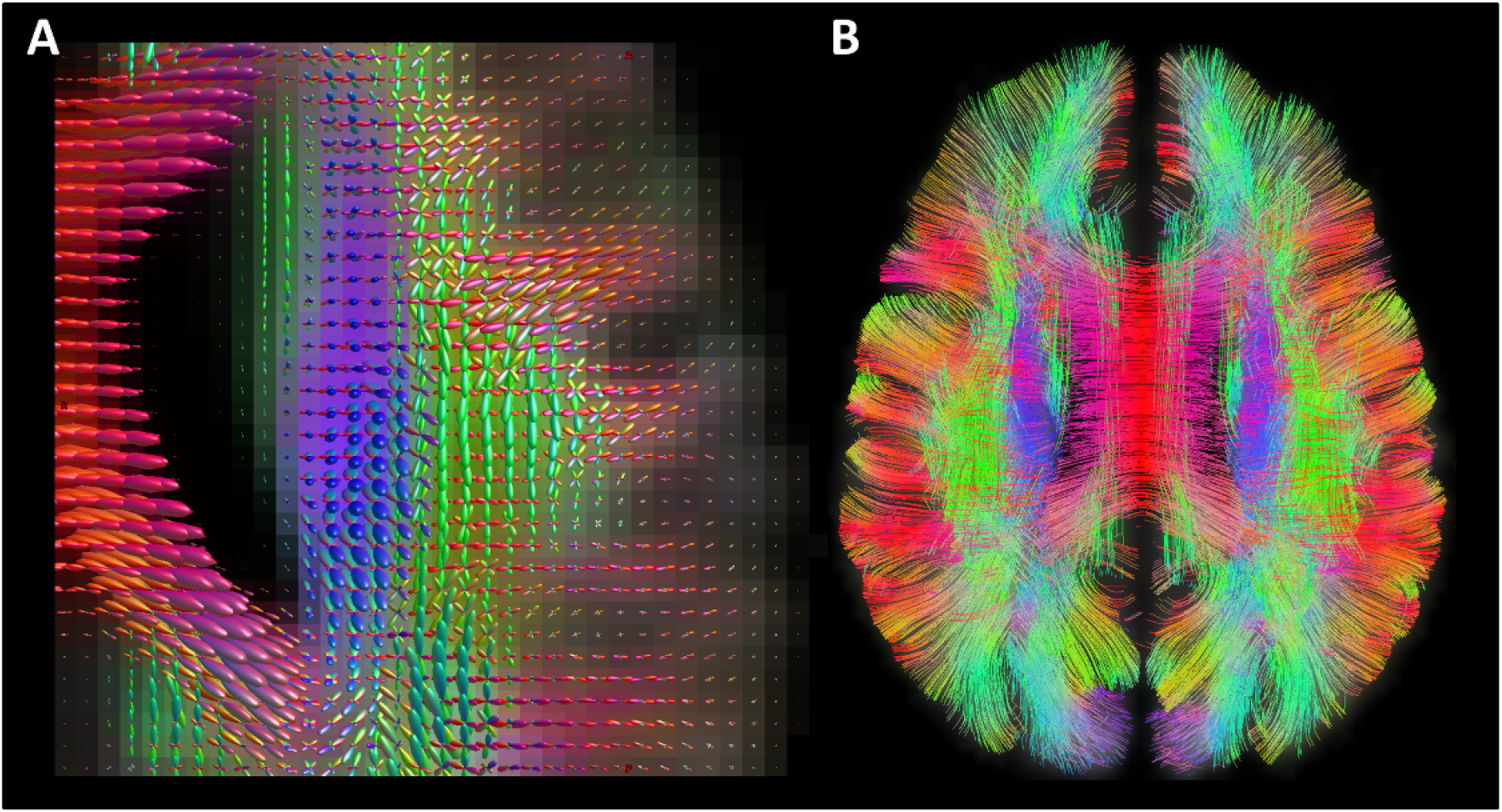
**A**. Visualization of fiber orientation distribution functions (fODFs) derived from the population template of MAP propagators shows multiple peaks in areas of crossing white matter fiber bundles. **B**. Whole-brain streamline fiber tractography using the template fODFs field can be used to resolve crossing white matter pathways.

Fig. 4 illustrates how the proposed framework for spatial standardization and normalization of MAPMRI data can be used to assess individual anatomical and microstructural differences between subjects in template space. Fig. 4A displays angular dissimilarity images between a subject and the template in multiple slices across the whole brain. These maps comprehensively quantify voxel-wise differences between the diffusion displacement profiles of each subject and the template with the highest values found generally near white matter boundaries and in cortical sulci and gyri. Fig. 4B illustrates the intersubject variability of the 3D MAP propagators using the same angular dissimilarity metric computed for multiple subjects in the same slice. The relatively low values indicate that after registration throughout the brain the diffusion propagators are similar across subjects. Propagator differences tend to be higher in the peripheral WM, where the intersubject variability is more pronounced due to individual anatomical differences in cortical folding patterns. The overall high degree of similarity, particularly within major WM fiber pathways and GM structures, gives hope that the diffusion propagator-based spatial standardization and analysis framework may detect complex and subtle microstructural changes during disease.

**Figure 4:**
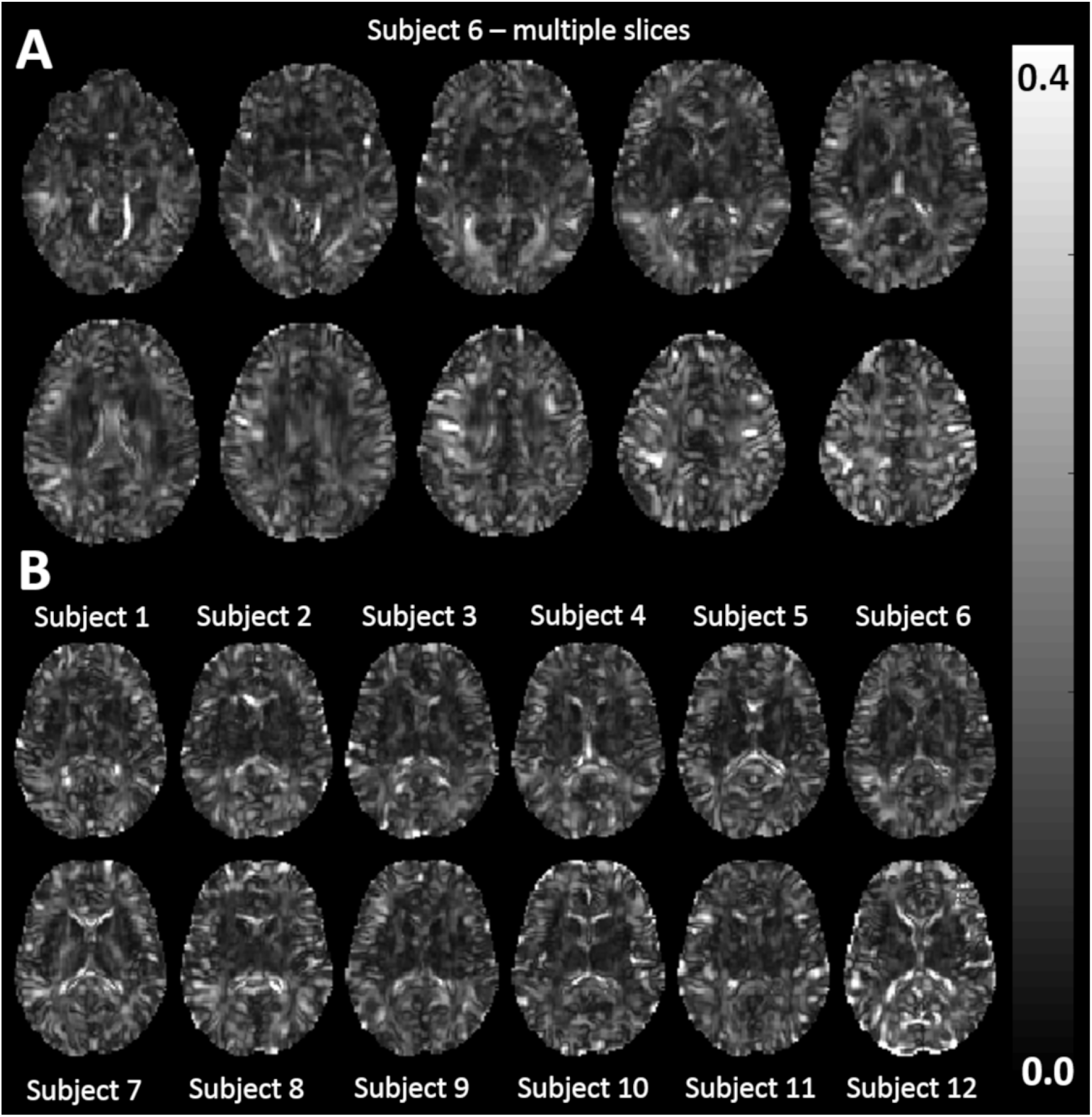
Voxelwise angular dissimilarity between single subject and template co-registered MAP-MRI propagators. **A**. Multiple slices from the same representative subject. **B**. The same representative slice for all subjects.

Figure 5 compares average MAP parameter values calculated in native and template space within each of the ICBM-81-DTI atlas ROIs. In native space, propagators are measured using a unique set of MAP basis functions computed from the subjects DTI data. In template space, propagators for all subjects are measured using the same set of voxelwise standardized MAP basis functions determined by the DTI population template. The average parameter value within each ROI across subjects in native space is plotted against the average parameter value within the same ROIs for across subjects in template space. A red line, with a slope of one is included for reference. The plots demonstrate that the NG provides the best agreement between the two types of ROI analyses. Values of RTOP and PA were relatively smaller in the subject-specific ROI-analysis. Such differences arise likely due to spatial interpolation during the transformation of MAP DWIs to template space. Acquiring MAP data with higher spatial resolution in future studies could mitigate these effects.

**Figure 5:**
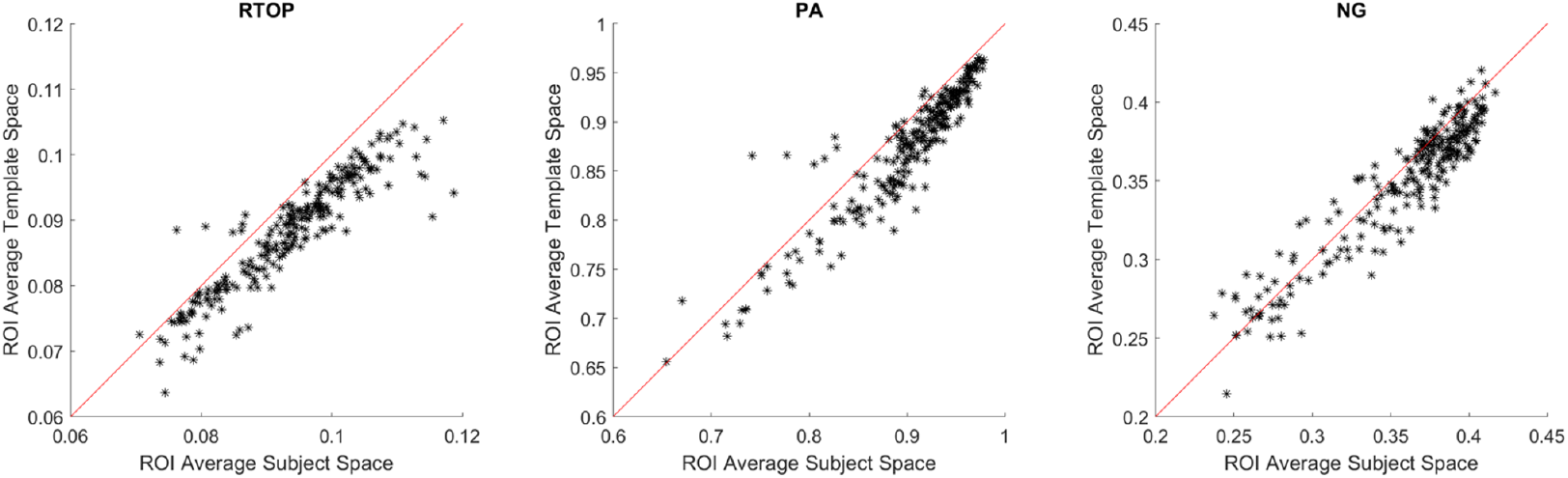
Correlation plots show good agreement between MAP-MRI derived parameters (RTOP left, PA middle, NG right) quantified using ROI-analyses in native and template space, respectively. In native space, propagators are measured using a unique set of MAP basis functions computed from each subjects DTI data. In template space, propagators for all subjects are measured using the same set of voxelwise standardized MAP basis functions determined by the DTI population template. In each plot, the x-axis corresponds to the ROI-average value of a MAP-MRI parameter in subject (native) space, while the y-axis corresponds to the ROI-average value of a MAP-MRI parameter in template space.

The data were further analyzed by examining the variability in ROI-averaged parameter values in all 50 WM ROIs across subjects using both techniques. Table 1 summarizes the average standard deviations of the estimated parameter values, and shows that the intersubject variability of PA and RTOP is higher when performing MAP analysis in subject space, while the NG shows similar intersubject variability in subject and template spaces.

**Table 1:**
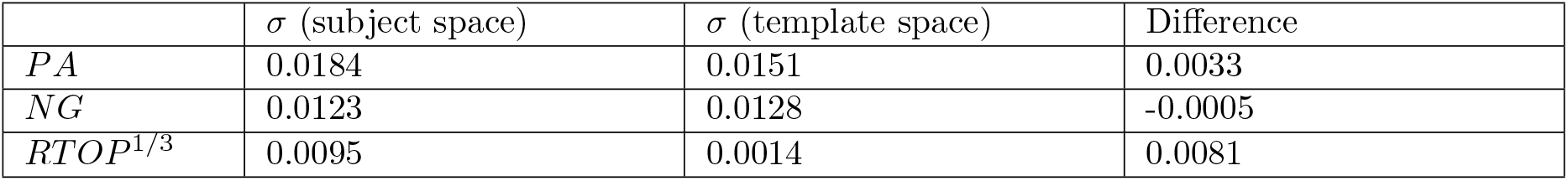
The intersubject standard deviation of the average PA (top row), NG (middle row) and RTOP (bottom row) across all 50 WM ROIs in subject (native) space (left) and template space (middle). The difference is shown in the last column.

| | $\sigma$ (subject space) | $\sigma$ (template space) | Difference |
| --- | --- | --- | --- |
| *PA* | 0.0184 | 0.0151 | 0.0033 |
| *NG* | 0.0123 | 0.0128 | -0.0005 |
| *RTOP<sup>1/3</sup>* | 0.0095 | 0.0014 | 0.0081 |

To illustrate how the proposed MAP-MRI analysis pipeline may be used clinically we analyzed longitudinal MAP data from a subject who underwent carotid endarterectomy for asymptomatic carotid atherosclerosis and was imaged at three timepoints: prior to the surgery, 1 month after surgery, and 6 months after surgery. The patient had no peri-operative or post-operative stroke during this period. The analysis did not find distinguishable differences between DTI or MAP microstructural parameters computed at the three timepoints in either native or template spaces. Angular dissimilarity values between propagators at different timepoints throughout most of the brain were comparable to those measured across propagators from multiple healthy subjects (Fig. 4). However, we observed localized differences between propagators at the three timepoints which stood out against the relatively low background angular dissimilarity and JSD values in the rest of the brain (Fig. 6). These very preliminary results call for a more extensive investigation to determine whether the observed changes are reproducible and related to the surgery in a larger cohort of patients. Future proof-of-principle and clinical validation studies will establish the sensitivity of the proposed MAP analysis framework for detecting local tissue changes longitudinally by quantifying propagator differences voxel-wise.

**Figure 6:**
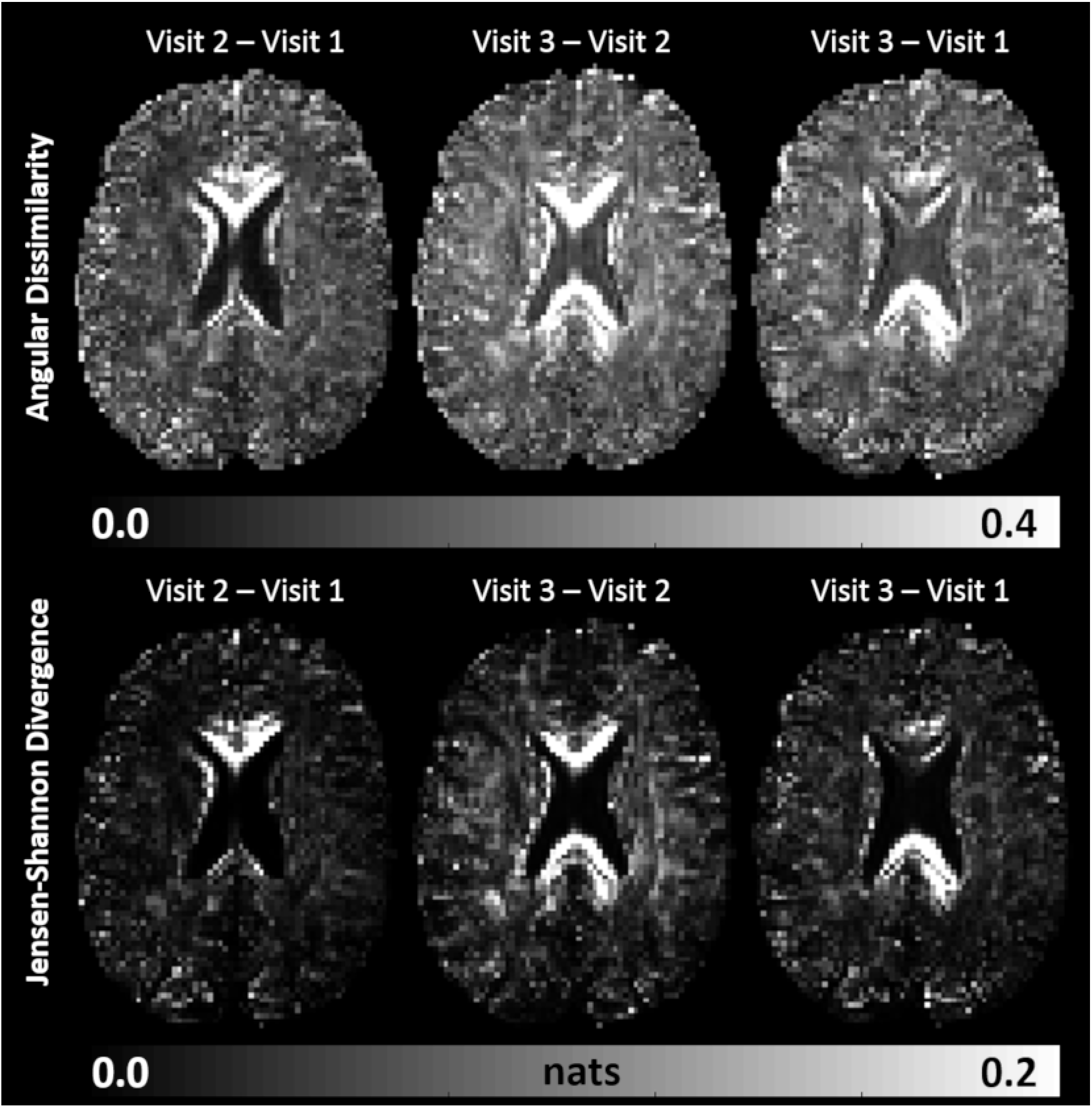
Quantifying voxelwise differences between MAP-MRI propagators acquired longitudinally in the same subject undergoing vascular occlusion surgery at three different clinical timepoints: Visit 1 - baseline. Visit 2 - 1 month after surgery. Visit 3 - 6 months after surgery. Maps of angular dissimilarity (top row) and Jensen-Shannon Divergence (bottom row) in the same representative slice.

## 4. Discussion

A good first approximation of the dMRI signal in both isotropic and anisotropic tissues can be obtained with DTI. MAP-MRI extends the DTI framework to explicitly measure diffusion propagators with arbitrary levels of complexity. Diffusion signals are represented in an orthogonal set of Gauss-Hermite basis functions constructed with orientation and scaling parameters derived from the principal directions and diffusivities of the diffusion tensor. The propagator is approximated efficiently by refining the tensor model with a linear combination of MAP basis functions (Appendix A). Since MAP-MRI subsumes and generalized DTI, it is well-suited for spatial standardization using tensor-based image registration algorithms, such as DRTAMAS (Irfanoglu et al., 2016). DRTAMAS provides morphologically accurate tensor-based registration by intelligently weighing the isotropic (trace) and anisotropic (deviatoric) components of the diffusion tensors during the alignment process.

Our results indicate that we can combine simple and well-established steps from existing tensor-based diffeomorphic registration pipelines (e.g., DRTAMAS) in order to register multiple MAP-MRI data sets for subsequent voxelwise analysis. Compared to ODF/fODF-based registration methods which align DWIs using spherical coordinates, tensor-based approaches may be less efficient in characterizing the orientational dependence of the propagators. Nevertheless, as shown in Fig. 3, our approach can readily distinguish crossing white matter pathways using whole-brain streamline fiber tractography while providing accurate quantitation of microstructural tissue parameters and propagator features (Fig. 5, Table 1). Moreover, unlike ODF/fODF-based methods, tensor-based diffeomorphic registration requires fewer optimization parameters, inherently works in Cartesian coordinates (compatible with DTI and MAP) and uses simple and well-established computational steps. Tensor-based dMRI registration pipelines are continuously tested and quality controlled in a growing number of applications, and have been shown to provide reliable results. The simplicity and legitimacy of each individual step is a strength of our approach.

Moreover, the use of tensor-based registration naturally provides a DTI study template. This diffusion tensor template is necessary in order to measure and analyze propagators using consistent sets of MAP basis functions across co-registered data sets. At each voxel, the MAP basis functions are derived from an average diffusion tensor (template DTI) that represents the best first-order approximation of the dMRI signals in all data sets. In some voxels prone to misregistration, propagators in template space may be approximated using suboptimal basis functions. However, we do not expect this to produce large quantitation errors. Due to the orthogonality of the MAP basis functions the propagator approximation generally converges efficiently even if the basis functions are not optimal (Avram et al., 2016).

On the other hand, analyzing co-registered propagators using MAP basis functions standardized for each location has several important advantages. The linearity of the MAP basis functions preserves important features of probability density functions after averaging multiple propagators, such as non-negativity and normalization (integrates to 1), and allows the direct averaging, analysis and comparison of propagators and their features using analytical formulas that can be very robust to measurement noise.

In general, the distinction between two propagators, can be assessed quantitatively using metrics other than the angular similarity defined in Eq. 1. Since diffusion propagators are probability density functions of net displacements of diffusing water molecules, they are well-suited for statistical quantitation using probability distance metrics. For example, a general measure of the difference in information content between two probability distributions is the Jensen-Shannon divergence (JSD) (Lin, 1991), a symmetrized extension of the KullbackLeibler divergence (KLD) between probability distributions (Kullback and Leibler, 1951). As shown in Fig. 6, the JSD and angular dissimilarity both highlight similar brain regions, with the JSD having a larger dynamic range, and potentially being more sensitive. Another advantage of computing the JSD is the possibility of comparing propagator differences to diffusion tensor differences measured using the KLD. Fig. 7 compares the relative entropy, i.e., KLD, derived from the DTI of a single subject and the DTI template to the total divergence, i.e., JSD, computed from the MAP data of the same subject and the MAP template. The relatively higher values of JSD compared to KLD reveal areas where the propagator-based analysis may be more sensitive to the individual anatomical or microstructural variations relative to the template, providing additional information compared to the DTI analysis. Probability distance measures between co-registered propagators could potentially reveal multifaceted microstructural changes that otherwise we could not detect solely based on microstructural DTI and scalar MAP parameters that assess specific features of the propagators such as size, anisotropy or non-gaussianity.

**Figure 7:**
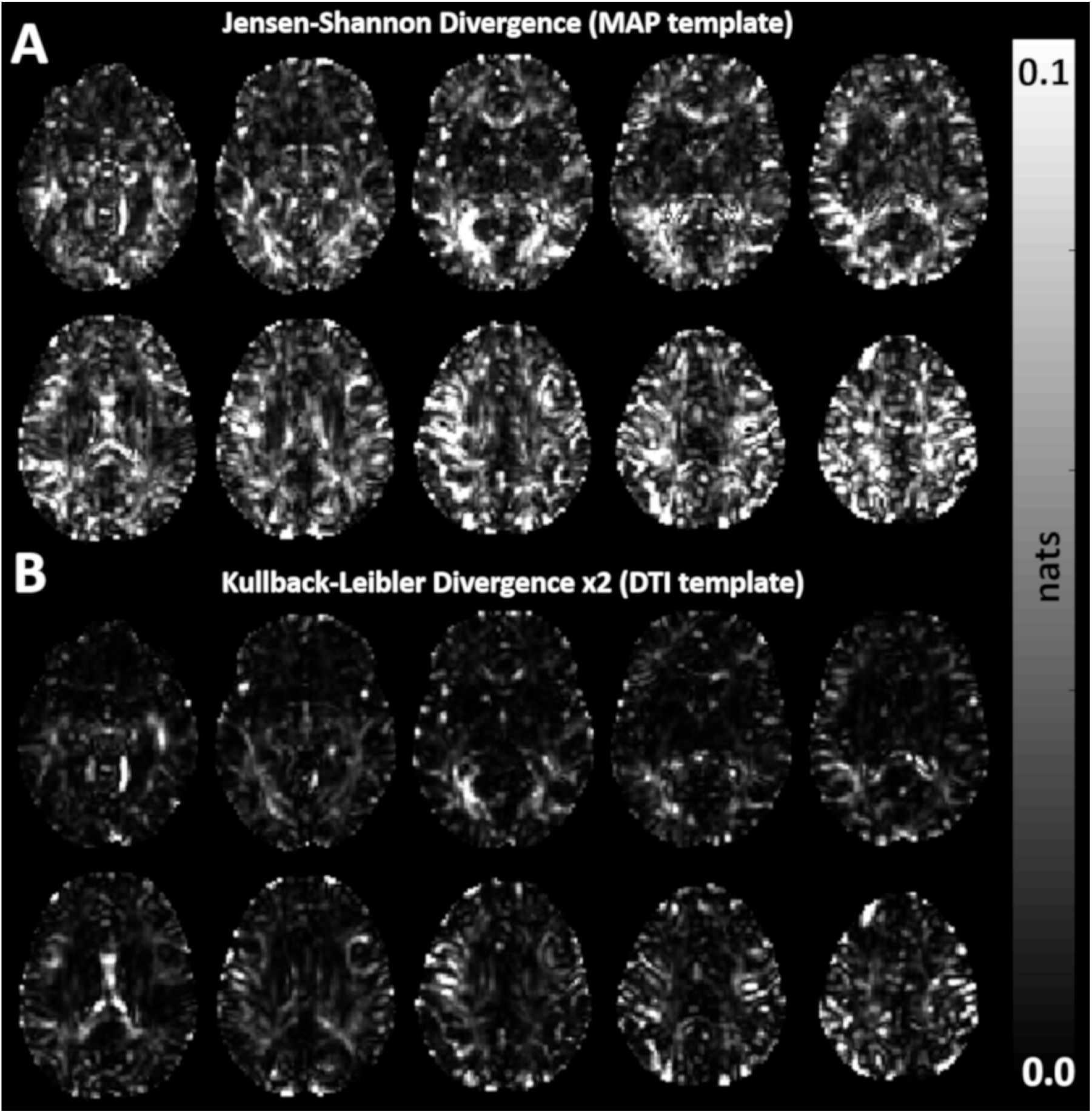
Probability distance measures (natural units of information) between the individual subject and template propagators measured with MAP (**A**) and diffusion tensors measured with DTI (**B**) in the same slices shown in Fig. 4A. The KullbakLeibler divergence (KLD) is used to quantify differences between diffusion tensors, while a symmetrized version of the KLD, called Jensen-Shannon divergence (JSD), is used to quantify differences between propagators. Note that the KLD is scaled by a factor of 2.

Voxelwise analysis of co-registered propagators may improve sensitivity in cross-sectional clinical studies. The intersubject variability of diffusion propagators within the cohort of healthy subjects was relatively low, with the largest angular dissimilarity values localized mainly in peripheral white matter regions. Our results in Fig. 5 indicate that the intersubject variability in the quantitation of MAP parameters is reduced, in general, by registering the MAP data to a common template as opposed to performing the analysis on a subject-by-subject basis. This is an important finding, as it suggests that the sensitivity of MAP-MRI parameters to changes in underlying microstructure may be increased by spatial standardization of MAP data sets in cross-sectional clinical studies. in general, the intersubject variability may be further reduced using tract-based spatial statistics (TBSS) (Smith et al., 2006). Nevertheless, TBSS analysis has numerous pitfalls associated with individual processing steps (Bach et al., 2014) that may obfuscate clinically relevant microstructural differences between data sets.

The registration framework from Fig. 1 may also help detect subtle longitudinal changes in microstructure during disease, treatment response, healthy development, or aging. Voxelwise comparison of propagators may be more eloquent and conspicuous in longitudinal MAP data sets in which anatomical structures with healthy tissues may show very little variability across timepoints, serving as internal control for the detection of fine pathophysiological alterations.

Finally, an interesting potential application of the proposed framework is the construction of improved anatomical brain atlases based on higher-order microstructural information (Varentsova et al., 2014) available in the MAP. Just as DTI parameters, such as the direction-encoded color (DEC) map provided a new contrast for delineating white matter fiber pathways in brain atlases (Mori et al., 2009), various propagators features in the MAP template may help improve delineation and visualization of anatomical brain structures. Using the entire diffusion propagators, including information about anisotropy, shape, size, ODFs, nongasussianty, zero-displacement probabilities, etc., may provide a very rich description of the tissue cytoarchitecture and microstructure and lead to more accurate and comprehensive MRI-based brain atlases. It will be particularly interesting to relate differences between the template propagators in white and gray matter to microstructural parameters derived with dMRI tissue models such as NODDI (Zhang et al., 2012). Future studies using larger MAP-MRI data sets with higher spatial resolution may enable us to identify cortical areas with similar distinct propagator features, indicative of microstructural and cytoarchitectural characteristics that correlate with postmortem histological stains.

## Acknowledgements

This work was supported by the Center for Neurodegeneration and Regenerative Medicine (CNRM), under the auspices of the Department of Defense (DoD) and the Henry Jackson Foundation (HJF) grant number #3080498.01-60855, and by the Intramural Research Program (IRP) of the National Institute of Biomedical Imaging and Bioengineering (NIBIB) and the *Eunice Kennedy Shriver* National Institute of Child Health and Human Development (NICHD) within the National Institutes of Health (NIH). We would also like to thank the University of Arizona for providing the longitudinal data set used in this study. The carotid endarterectomy work was supported by the University of Arizona Translational Imaging Program Project Stimulus (TIPPS) grant.

## A. Appendix A: The MAP-MRI basis functions

Mean apparent propagator (MAP) MRI (Özarslan et al., 2013) measures the probability density function of net 3D displacements of diffusing water molecules, i.e., the apparent diffusion propagator *P* (**r**), by analyzing multiple measurements in q-space, **q** = (2*π*)^*−*1^ *γδ***G**, with a series approximation using a complete set of orthogonal 3D Gauss-Hermite basis functions. The MAP-MRI basis functions are defined in the frame of reference that diagonalizes the diffusion tensor, **D**, using spatial scaling parameters derived from the diffusion time, *t*_*d*_, and the eigenvalues of the diffusion tensor, *λ*_*i*_, as 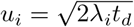, for *i* = *x, y, z*.

To measure the vector of MAP-MRI coefficients, 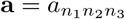, we first rotate each q-vector to the reference frame that diagonalizes the diffusion tensor, then fit the diffusion signal *E* (**q**) with the vector of MAP basis functions 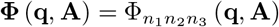 defined in the DTI reference frame using the diagonal 3 *×* 3 scaling matrix **A** = 2**D***t*_*d*_:

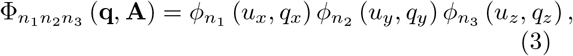

where

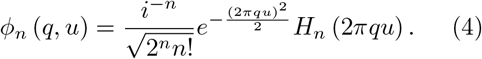

Each of the separable functions 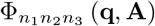 is determined by the spatial scaling coefficients *u*_*x*_, *u*_*y*_, and *u*_*z*_, and by the indices *n*_1_, *n*_2_, and *n*_3_, respectively corresponding to orders of the physicist’s Hermite polynomials *H* (*x*) in the orthogonal spatial dimensions of the DTI reference frame.

Since the Gauss-Hermite functions in MAPMRI are eigenfunctions of the Fourier Transform, the same MAP coefficients, **a**, describe both *E* (**q**, **A**) = **a^T^Φ** (**q**, **A**), and its Fourier Transform, *P* (**r**, **A**) = **a**^**T**^**Ψ** (**r**, **A**), where the vector 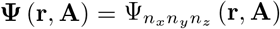 contains the MAP basis functions

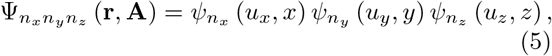

and

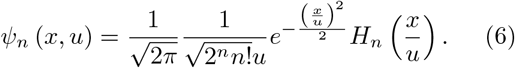

## B Appendix B: Approximation of the Jensen-Shannon divergence for MAP propagators

The information content for a probability density function can be quantified using the Shannon entropy (self-information) measure from information theory:

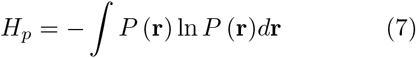

The Kullback-Leibler divergence, also known as the information divergence, information gain, or relative entropy, is a convenient way to compare two probability density functions P and Q, and is defined as:

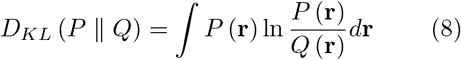

*D*_*KL*_ divergence is always non-negative, remains well-defined for continuous distributions and is invariant under parameter transformation. Moreover, the KL divergence is additive for independent distributions.

The Jensen-Shannon divergence is an extension of *D*_*KL*_ and has additional useful properties. It is symmetric and always finite and therefore can be used as a mutual information (or difference) metric between two probability distributions. For two continuous distributions *P* (**r**) and *Q* (**r**), it is defined as:

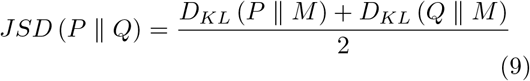

where 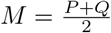.

To compute analytically an estimate of the JSD for two MAP propagators, we can use Eq. 3 to approximate ln *P* (**r**) as:

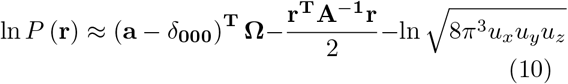

In the expression above *δ*_**000**_ is a column vector with a value of 1 corresponding to the element with indices *n*_1_ = *n*_2_ = *n*_3_ = 0, and zeros everywhere else. We have used the approximation ln (1 + *x*) ≈ *x* for small values of *x*, and have made the additional assumption that the first term of the MAP series expansion, *a*_000_, corresponding to the diffusion tensor component is the dominant term and *a*_000_ ≈ 1, and explains most of the diffusion MRI signal. This is a reasonable approximation especially for *in vivo* MAP-MRI data (Avram et al., 2016).

For two propagators, *P* (**r**) and *Q* (**r**), defined by MAP coefficient column vectors **a** and **b**, respectively, in the same MAP basis function determined by scaling parameters *u*_*x*_, *u*_*y*_, and *u*_*z*_, we can use Eq. 6–8 to obtain:

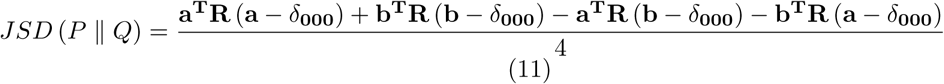

where **R** = ∫ **Ψ**^**T**^**Ω***d***r** is a constant matrix whose elements, for 3D MAP-MRI, are given by the product 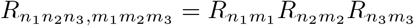 (Avram et al., 2017), with

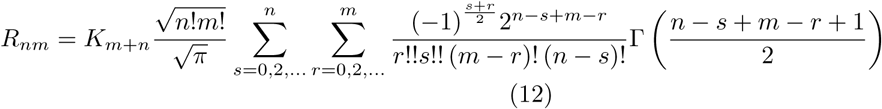

*K*_*m*+*n*_ is non-zero only if m and n are non-zero. Γ (*x*) represents the complete Gamma function. Note that that if we want to evaluate the bounded integral 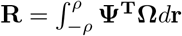 in order to mitigate the effects from oscillations of the MAP functions at larger displacements, we can simply replace the complete gamma function Γ (*x*) with the incomplete gamma function *γ* (*x*; *ρ*) in Eq. 10.

## References

Alexander, D. C., Pierpaoli, C., Basser, P. J., Gee, J. C., 2001. Spatial transformations of diffusion tensor magnetic resonance images. IEEE Transactions on Medical Imaging 20 (11), 1131–1139.

Avram, A., Hutchinson, E., Basser, P., 2017. Higher-order statistics of 3D spin displacement probability distributions measured with MAP MRI. In: Proceedings of the ISMRM. Vol. 25. p. 3367.

Avram, A. V., Barnett, A., Basser, P., 2014. The variation of MAP-MRI derived parameters along white matter fiber pathways in the human brain. In: Proceedings of the ISMRM. Vol. 22. p. 2587.

Avram, A. V., Sarlls, J. E., Barnett, A. S., zarslan, E., Thomas, C., Irfanoglu, M. O., Hutchinson, E., Pierpaoli, C., Basser, P. J., 2016. Clinical feasibility of using mean apparent propagator (MAP) MRI to characterize brain tissue microstructure. NeuroImage 127, 422–434.

Bach, M., Laun, F. B., Leemans, A., Tax, C. M. W., Biessels, G. J., Stieltjes, B., Maier-Hein, K. H., 2014. Methodological considerations on tract-based spatial statistics (tbss). NeuroImage 100, 358–369.

Barmpoutis, A., Vemuri, B. C., Forder, J. R., 2007. Registration of high angular resolution diffusion mri images using 4 th order tensors. In: International Conference on Medical Image Computing and Computer-Assisted Intervention. Springer, pp. 908–915.

Basser, P. J., Mattiello, J., LeBihan, D., 1994. MR diffusion tensor spectroscopy and imaging. Biophysical journal 66 (1), 259–67.

Caruyer, E., Lenglet, C., Sapiro, G., Deriche, R., 2013. Design of multishell sampling schemes with uniform coverage in diffusion MRI. Magnetic Resonance in Medicine 69 (6), 1534–1540.

Cheng, J., Shen, D., Yap, P., 2014. Designing single- and multiple-shell sampling schemes for diffusion mri using spherical code. In: Medical Image Computing and Computer-Assisted Intervention, MICCAI 2014 - 17th International Conference, Proceedings, 3rd Edition. Vol. 8675 LNCS. Springer Verlag, Germany, pp. 281–288.

Chiang, M.-C., Leow, A. D., Klunder, A. D., Dutton, R. A., Barysheva, M., Rose, S. E., McMahon, K. L., De Zubicaray, G. I., Toga, A. W., Thompson, P. M., 2008. Fluid registration of diffusion tensor images using information theory. IEEE transactions on medical imaging 27 (4), 442–456.

Garaci, F., Toschi, N., Lanzafame, S., Marfia, G. A., Marziali, S., Meschini, A., Di Giuliano, F., Simonetti, G., Guerrisi, M., Massa, R., Floris, R., 2015. Brain MR diffusion tensor imaging in Kennedy’s disease. Neuroradiology Journal 28 (2), 126–132.

Gee, J. C., Alexander, D. C., 2006. Diffusion-tensor image registration. In: Weickert, J., Hagen, H. (Eds.), Visualization and Processing of Tensor Fields. Springer Berlin Heidelberg, Berlin, Heidelberg, pp. 327–342.

Geng, X., Ross, T. J., Zhan, W., Gu, H., Chao, Y.-P., Lin, C.-P., Christensen, G. E., Schuff, N., Yang, Y., 2009. Diffusion mri registration using orientation distribution functions. In: International Conference on Information Processing in Medical Imaging. Springer, pp. 626–637.

Ginsburger, K., Poupon, F., Teillac, A., Mangin, J.-F., Poupon, C., 2018. Diffeomorphic registration of diffusion mean apparent propagator fields using dynamic programming on a minimum spanning tree. In: Kaden, E., Grussu, F., Ning, L., Tax, C. M. W., Veraart, J. (Eds.), Computational Diffusion MRI. Springer International Publishing, pp. 81–90.

Irfanoglu, M. O., Modi, P., Nayak, A., Hutchinson, E. B., Sarlls, J., Pierpaoli, C., 2015. DR-BUDDI: (Diffeomorphic registration for blip-up blip-down diffusion imaging) method for correcting echo planar imaging distortions. Neuroimage 106, 284–289.

Irfanoglu, M. O., Nayak, A., Jenkins, J., Hutchinson, E. B., Sadeghi, N., Thomas, C. P., Pierpaoli, C., 2016. DRTAMAS: Diffeomorphic registration for tensor accurate alignment of anatomical structures. NeuroImage 132, 439–454.

Jones, D. K., Griffin, L. D., Alexander, D. C., Catani, M., Horsfield, M. A., Howard, R., Williams, S. C. R., 2002. Spatial normalization and averaging of diffusion tensor MRI data sets. NeuroImage 17 (2), 592–617.

Kellner, E., Dhital, B., Kiselev, V. G., Reisert, M., 2016. Gibbs-ringing artifact removal based on local subvoxel-shifts. Magnetic Resonance in Medicine 76 (5), 1574–1581.

Kullback, S., Leibler, R. A., 03 1951. On information and sufficiency. The Annals of Mathematical Statistics 22 (1), 79–86.

Lin, J., Jan. 1991. Divergence measures based on the Shannon entropy. IEEE Transactions on Information Theory 37 (1), 145–151.

Mahoney, C. J., Simpson, I. J. A., Nicholas, J. M., Fletcher, P. D., Downey, L. E., Golden, H. L., Clark, C. N., Schmitz, N., Rohrer, J. D., Schott, J. M., Zhang, H., Ourselin, S., Warren, J. D., Fox, N. C., 2015. Longitudinal diffusion tensor imaging in rontotemporal dementia. Annals of Neurology 77 (1), 33–46.

Maier-Hein, K. H., Westin, C., Shenton, M. E., Weiner, M. W., Raj, A., Thomann, P., Kikinis, R., Stieltjes, B., Pasternak, O., 2015. Widespread white matter degeneration preceding the onset of dementia. Alzheimer’s and Dementia 11 (5), 485–493.

Mori, S., Oishi, K., Faria, A. V., 2009. White matter atlases based on diffusion tensor imaging. Current Opinion in Neurology 22 (4), 362–369.

Muoz-Moreno, E., Crdenes-Almeida, R., Martin-Fernandez, M., 2009. Review of techniques for registration of diffusion tensor imaging. Advances in Pattern Recognition, 273–297.

Özarslan, E., Koay, C. G., Shepherd, T. M., Komlosh, M. E., rfanolu, M. O., Pierpaoli, C., Basser, P. J., 2013. Mean apparent propagator (MAP) MRI: A novel diffusion imaging method for mapping tissue microstructure. NeuroImage 78 (0), 16–32.

Pierpaoli, C., Walker, L., Irfanoglu, M., Barnett, A., Basser, P., Chang, L., Koay, C., Pajevic, S., Rohde, G., Sarlls, J., 2010. TORTOISE: an integrated software package for processing of diffusion MRI data. In: Proceedings of the ISMRM. Vol. 18. p. 1597.

Poudel, G. R., Stout, J. C., Domnguez D., J. F., Churchyard, A., Chua, P., Egan, G. F., Georgiou-Karistianis, N., 2015. Longitudinal change in white matter microstructure in Huntington’s disease: The IMAGE-HD study. Neurobiology of disease 74 (1), 406–412.

Raffelt, D., Tournier, J. D., Fripp, J., Crozier, S., Connelly, A., Salvado, O., 2011. Symmetric diffeomorphic registration of fibre orientation distributions. NeuroImage 56 (3), 1171–1180.

Sadeghi, N., Nayak, A., Walker, L., Okan Irfanoglu, M., Albert, P. S., Pierpaoli, C., Group, B. D. C., 2015. Analysis of the contribution of experimental bias, experimental noise, and inter-subject biological variability on the assessment of developmental trajectories in diffusion MRI studies of the brain. NeuroImage 109, 480–492.

Smith, S. M., Jenkinson, M., Johansen-Berg, H., Rueckert, D., Nichols, T. E., Mackay, C. E., Watkins, K. E., Ciccarelli, O., Cader, M. Z., Matthews, P. M., Behrens, T. E. J., 2006. Tract-based spatial statistics: Voxelwise analysis of multi-subject diffusion data. NeuroImage 31 (4), 1487–1505.

Tournier, J. D., Calamante, F., Connelly, A., 2007. Robust determination of the fibre orientation distribution in diffusion mri: Non-negativity constrained super-resolved spherical deconvolution. NeuroImage 35 (4), 1459–1472.

Tournier, J.-D., Calamante, F., Connelly, A., 2012. Mrtrix: Diffusion tractography in crossing fiber regions. International Journal of Imaging Systems and Technology 22 (1), 53–66.

Tournier, J.-D., Calamante, F., Gadian, D. G., Connelly, A., 2004. Direct estimation of the fiber orientation density function from diffusion-weighted mri data using spherical deconvolution. NeuroImage 23 (3), 1176 – 1185.

Varentsova, A., Zhang, S., Arfanakis, K., 2014. Development of a high angular resolution diffusion imaging human brain template. NeuroImage 91, 177–186.

Veraart, J., Fieremans, E., Novikov, D. S., nov 2015. Diffusion MRI noise mapping using random matrix theory. Magnetic Resonance in Medicine 76 (5), 1582–1593.

Wang, Y., Gupta, A., Liu, Z., Zhang, H., Escolar, M. L., Gilmore, J. H., Gouttard, S., Fillard, P., Maltbie, E., Gerig, G., Styner, M., 2011. DTI registration in atlas based fiber analysis of infantile Krabbe disease. NeuroImage 55 (4), 1577–1586.

Watts, R., Thomas, A., Filippi, C. G., Nickerson, J. P., Freeman, K., 2014. Potholes and molehills: Bias in the diagnostic performance of diffusion-tensor imaging in concussion. Radiology 272 (1), 217–223.

White, T., Schmidt, M., Karatekin, C., 2009. White matter potholes in early-onset schizophrenia: a new approach to evaluate white matter microstructure using diffusion tensor imaging. Psychiatry research 174 (2), 110–115.

Zhang, H., Schneider, T., Wheeler-Kingshott, C. A., Alexander, D. C., 2012. Noddi: Practical in vivo neurite orientation dispersion and density imaging of the human brain. NeuroImage 61 (4), 1000–1016.

Zhang, H., Yushkevich, P. A., Alexander, D. C., Gee, J. C., 2006. Deformable registration of diffusion tensor MR images with explicit orientation optimization. Medical Image Analysis 10 (5), 764–785.

Zhang, P., Niethammer, M., Shen, D., Yap, P.-T., 2014. Large deformation diffeomorphic registration of diffusion-weighted imaging data. Medical Image Analysis 18 (8), 1290–1298.

Zhang, S., Arfanakis, K., 2018. Evaluation of standardized and study-specific diffusion tensor imaging templates of the adult human brain: Template characteristics, spatial normalization accuracy, and detection of small inter-group FA differences. NeuroImage 172, 40–50.

